# Mechanical contributions of the Myp2 tail revealed by coiled-coil force sensors

**DOI:** 10.64898/2026.06.18.733274

**Authors:** Juhyeong Lee, Julien Berro, Shinji Deguchi, Takumi Saito

## Abstract

Cytokinesis is driven by the constriction of an actomyosin contractile ring, yet how forces generated by individual myosin molecules are transmitted within the ring remains poorly understood. In fission yeast *Schizosaccharomyces pombe*, the type II myosin Myp2 plays a key role in ring constriction and stability, particularly under stress conditions. Unlike typical type-II myosins, Myp2’s tail contains two coiled coils. Although numerous studies have revealed the structure and molecular function of Myp2, the mechanical contribution of its unique tail region has remained unclear. Here, we directly measured forces acting on Myp2 in living cells using genetically encoded coiled-coil force sensors. Insertion of force sensors into the neck region revealed that Myp2 experiences forces of approximately 6.6 pN within the contractile ring. To dissect the contribution of the tail region, we systematically analyzed tail-deletion mutants. Deletion of the proximal tail had little effect on force levels, whereas deletion of the distal C-terminal tail significantly reduced the forces experienced by Myp2. Quantitative analysis of force distributions further showed that truncation of the C-terminal tail reduces the characteristic force by approximately half, demonstrating a major mechanical contribution of the distal tail to the forces experienced by Myp2 at the molecular scale. Deletion of the C-terminal tail slowed the growth rate, consistent with previous macroscopic observations that this region is required for robust ring constriction under stress conditions. We further found that the C-terminal tail forms clusters, which require full-length Myp2 for proper localization to the ring. These findings, together with molecular force measurements and quantitative analysis, demonstrate that the Myp2 tail is not merely a structural or regulatory domain but functions as a load-bearing component *in vivo*, with its C-terminal region playing a central role in force transmission across scales.

## Introduction

Cytokinesis is the final step of the cell cycle, during which a single cell physically divides into two daughter cells. In most eukaryotes, cytokinesis is driven by the assembly and constriction of an actomyosin contractile ring. This contractile ring generates mechanical forces that deform the plasma membrane and underlying cortex, leading to cleavage furrow ingression and abscission ^1,2^. Actin filaments and myosin-II motors cooperate to organize, regulate, and transmit forces across length scales, from molecular interactions to whole-cell mechanics ^3^.

While the basic architecture and molecules of the contractile ring are conserved among fungi and animals, the mechanisms by which molecular forces generated by myosin motors are coordinated and transmitted to produce robust and symmetric constriction remain incompletely understood. In animal cells, the contractile ring is embedded in a complex cortical network and is regulated by RhoA-mediated signaling pathways that modulate actin assembly, myosin activation, and filament crosslinking ^4,5^. Despite extensive characterization of the molecular components of cytokinesis, direct measurements of forces within contractile ring proteins have been technically challenging *in vivo*.

Fission yeast *Schizosaccharomyces pombe* has been used as a powerful model system for dissecting the molecular and mechanical basis of cytokinesis. Owing to its relatively simple geometry and genetic tractability, and the presence of highly conserved cytokinetic proteins, fission yeast allows quantitative studies that connect molecular interactions to contractile ring dynamics ^2,6^. Numerous studies combined with genetic manipulations, imaging techniques primarily relying on super resolution microscopy, and theoretical approaches have revealed the nanoscale architecture and molecular organization as well as the mechanical behavior of the ring ^7–10^.

Fission yeast expresses two type-II myosins, Myo2 and Myp2 (also known as Myo3), in cytokinesis. Myo2 is essential and plays a central role during early stages of contractile ring assembly and constriction, whereas Myp2 is non-essential but contributes significantly to ring stability and efficient constriction, particularly under stress conditions ^11–14^. Unlike canonical type-II myosins, Myp2 possesses a unique tail composed of two coiled-coil regions connected by a nonhelical hinge, forming an antiparallel intramolecular coiled coil and behaving as a monomeric, single-headed myosin ^15^ (Fig. 1A). This unusual tail structure, particularly its C-terminal distal region, is required for efficient cytokinesis and robust contractile-ring constriction *in vivo* ^15^. Genetic and cell-scale studies further suggest that Myp2 contributes to maintaining ring function under conditions of elevated mechanical or environmental stress ^14^.

**Figure 1.**
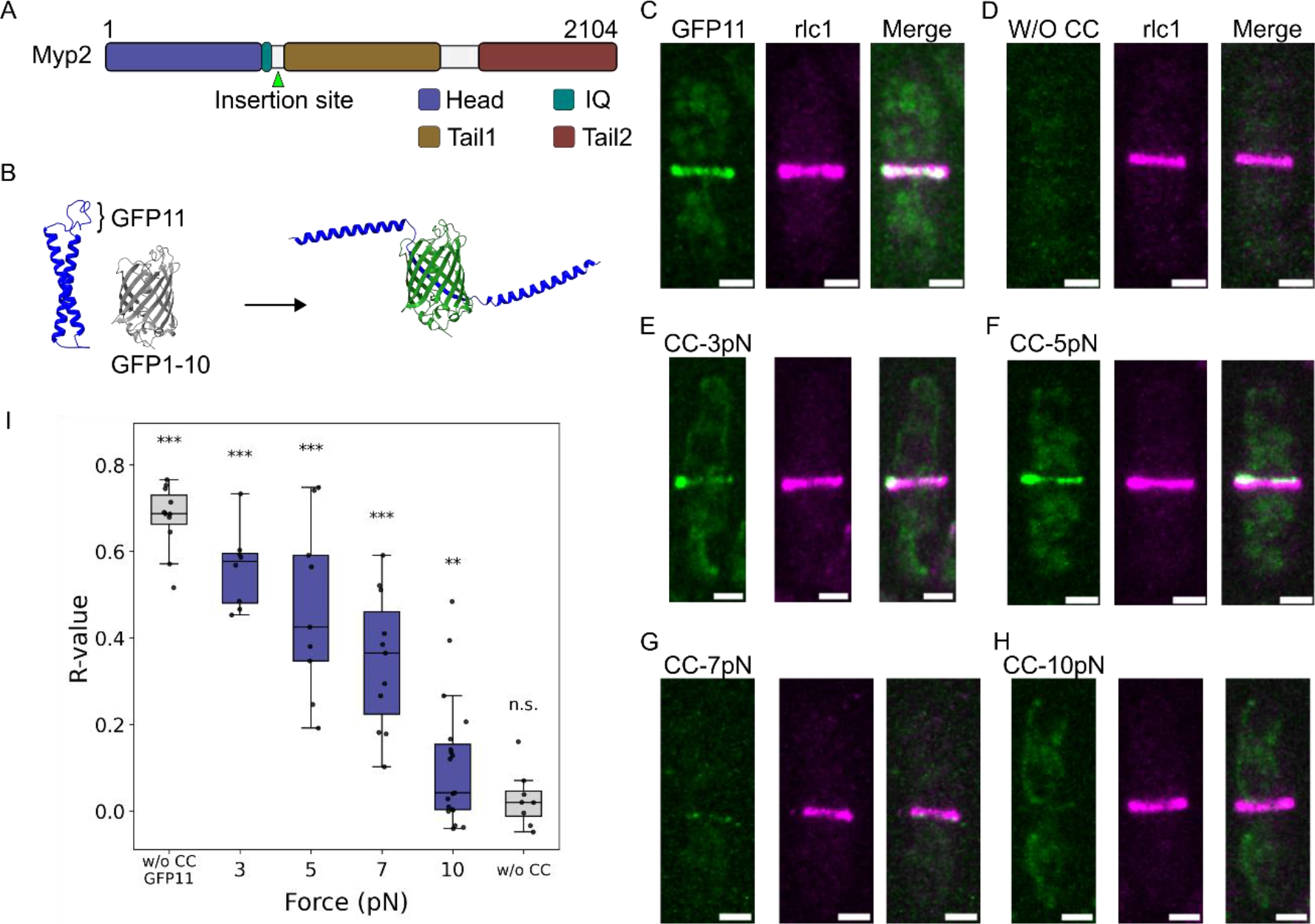
Force measurements on fission yeast type-II myosin Myp2 using coiled-coil (CC) force sensors. (A) Domain organization of Myp2. (B) Schematic illustration of a CC force sensor with split-GFP readout, predicted by AlphaFold 3 ^41^ visualized by UCSF ChimeraX ^42^.(C-H) Fluorescence images of strains expressing Rlc1-mScarlet3 and Myp2 with the indicated insertions. (C) GFP11 without CC force sensors. (D) No insertion. (E-H) CC force sensors with thresholds of (E) 3 pN, (F) 5 pN, (G) 7 pN, and (H) 10 pN. (I) Boxplots of colocalization analysis between GFP and Rlc1-mScarlet3, quantified by Pearson’s correlation coefficient (R-value). Statistical significances were determined by Permutation test with randomized intensity distribution. ***, p < 0.001; **, p < 0.01; and n.s., no significance. n > 8 cells per genotype. Scale bars; 2 μm.

Although this unique structure of the Myp2 tail has been extensively characterized both *in vitro* and *in vivo*, and genetic studies have revealed its importance for maintaining contractile ring function under stress conditions, the mechanical significance of whether Myp2 and its unique tail region directly contribute to mechanical forces at the molecular scale remains unclear. Here, we address this gap by directly measuring forces on Myp2 and its tail-deletion mutants in living cells using our genetically encoded coiled-coil force sensors ^16^.

## Results

### Myp2 bears more than 7 pN forces in contractile rings

The coiled-coil (CC) force sensor with split-GFP readout used in this study is composed of two helices and 11^th^ beta sheet of GFP as the linker ^16,17^. Once forces open the CC structure and expose the GFP11 linker, cytoplasmic diffusive GFP1-10 binds to the linker, forming a reconstituted whole GFP structure ^16,18^, meaning that fluorescent GFP signal indicate that CC force sensors are at the open state (Fig. 1B). Because the CC sensor is irreversible and lacks temporal resolution, the measured signal reflects cumulative force exposure rather than instantaneous force. By using CC force sensors with distinct force thresholds, we obtain a range of forces on a protein of interest.

To measure forces on Myp2, we genetically inserted CC sensors with spilt-GFP readout into Q833, the neck region between the IQ motif and tail1 in the strain expressing GFP1-10. Rlc1 was tagged with mScarlet3 as a contractile ring marker. GFP11 without CC was inserted into the insertion site as a positive control (Fig. 1C), whereas a strain without any insertion was used to obtain the background GFP signal in the ring (Fig. 1D). CC force sensors with distinct force thresholds from 3 pN to 10 pN were inserted at the same insertion site (Fig. 1E-H, S2). CC-3pN, CC-5pN, and CC-7pN sensors were open and fluorescent in contractile rings due to exposure of the GFP11 linker (Fig. 1E-G, S2), whereas CC-10pN showed weak fluorescence that was difficult to distinguish from background (Fig. 1H).

To quantify the levels of GFP signal due to Myp2 in the ring, colocalization analysis between Rlc1 and GFP was conducted at the single cell level (Fig. 1I). Strains expressing CC-3pN, CC-5pN, and CC-7pN exhibited statistically significant Pearson’s correlation coefficient (R-value) ranging from 0.3 to 0.5. Colocalization analysis of CC-10pN yielded a statistically significant but low R-value (R ≈ 0.1), indicating partial opening of the CC-10pN sensor (Fig. 1I). Taken together, these results indicate Myp2 bears forces up to values between 7 pN and 10 pN.

### Myp2 C-terminal distal tail is required for efficient force bearing by Myp2

Compared with other non-muscle type-II myosins, Myp2 has a unique domain organization composed of two distinct coiled-coil regions (hereafter called tail1 and tail2) connected by a non-coiled hinge ^15^. This unusual organization is important to support stable and robust ring constriction ^11,14^. Here, we asked whether these distinct tail regions contribute to forces on Myp2 at the molecular scale.

We first constructed a tail1 deletion mutant, Myp2-Δtail1 and inserted CC sensors at the same position as in full-length Myp2 in a strain expressing GFP1-10 (Fig. 2A). Similar to full-length Myp2, CC-3pN, CC-5pN, and CC-7pN sensors were predominantly open and fluorescent (Fig. 2B-D), whereas CC-10pN exhibited very weak fluorescence in contractile rings (Fig. 2E). Colocalization analysis indicated that the deletion of tail1 did not drastically alter force levels compared with full-length Myp2 (Fig. 2F).

**Figure 2.**
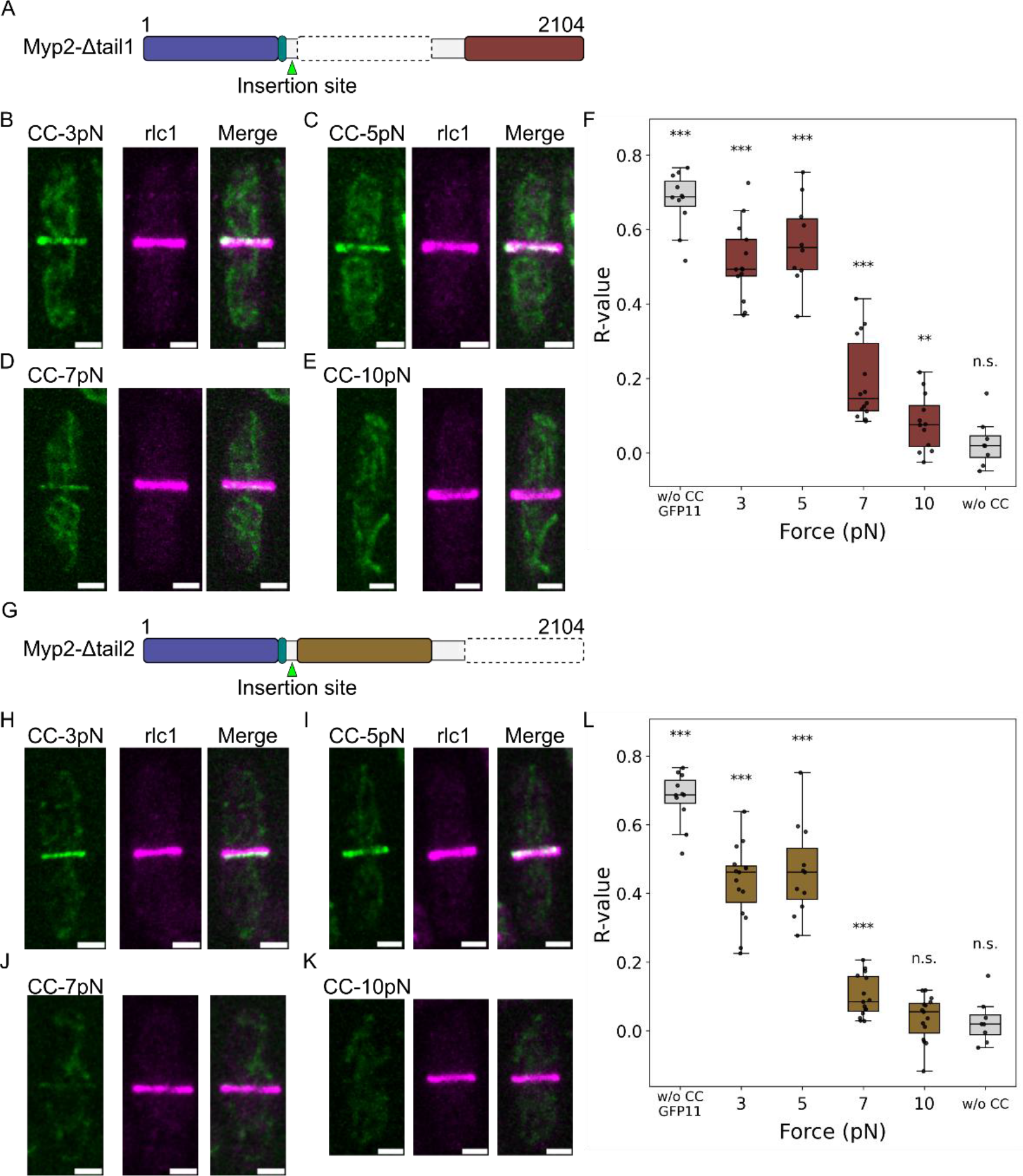
Force measurements on Myp2 deletion mutants using CC force sensors. (A) Schematic representation of the domain organization of the Myp2 proximal tail deletion mutant (Myp2-Δtail1). (B-E) Fluorescence images of strains expressing Rlc1-mScarlet3 and Myp2-Δtail1 with the indicated CC force sensors. (F) Boxplots of colocalization analysis between GFP and Rlc1-mScarlet3 for the Myp2-Δtail1 mutants. n >… cells per genotype. (G) Schematic representation of the domain organization of the Myp2 distal tail deletion (Myp2- Δ tail2). (H-K) Fluorescence images of strains expressing Rlc1-mScarlet3 and Myp2- Δ tail2 with the indicated CC force sensors. (L) Boxplots of colocalization analysis between GFP and Rlc1-mScarlet3 for the Myp2-Δtail2 mutants. Statistical significances were determined by Permutation test with randomized intensity distribution. n > 8 cells per genotype. ***, p < 0.001; **, p < 0.01; and n.s., no significance. Scale bars; 2 μm.

Next, a tail2 deletion mutant, Myp2-Δtail2, was constructed, and CC sensors were again inserted into the neck region (Fig. 2G). While fluorescence signals from CC-3pN, CC-5pN, and CC-7pN were observed (Fig. 2H-J), CC-7pN showed reduced GFP intensity and lower R-value (Fig. 2L) compared with full-length Myp2 and Myp2-Δtail1 (Fig. 1G, 1I, 2D, 2F). The GFP signal from CC-10pN was difficult to distinguish from background (Fig. 2K), resulting in a non-statistically significant R-value (Fig. 2L).

### Force measurements on Myp2 quantify mechanical significance of Myp2 tail regions

Because CC force sensors with the split-GFP readout detect the maximum force on a probing protein in a time-irreversible manner, GFP intensity can be used as proxy for the proportion of Myp2 molecules that experienced forces exceeding a given threshold (*θ*_cc_).

Thus, we analyzed GFP intensity in contractile rings for each genotype, revealing a monotonic decrease in GFP intensity with increasing CC force threshold (Fig. 3A-D). This result is consistent with a reduced probability of CC opening at higher force thresholds.

**Figure 3.**
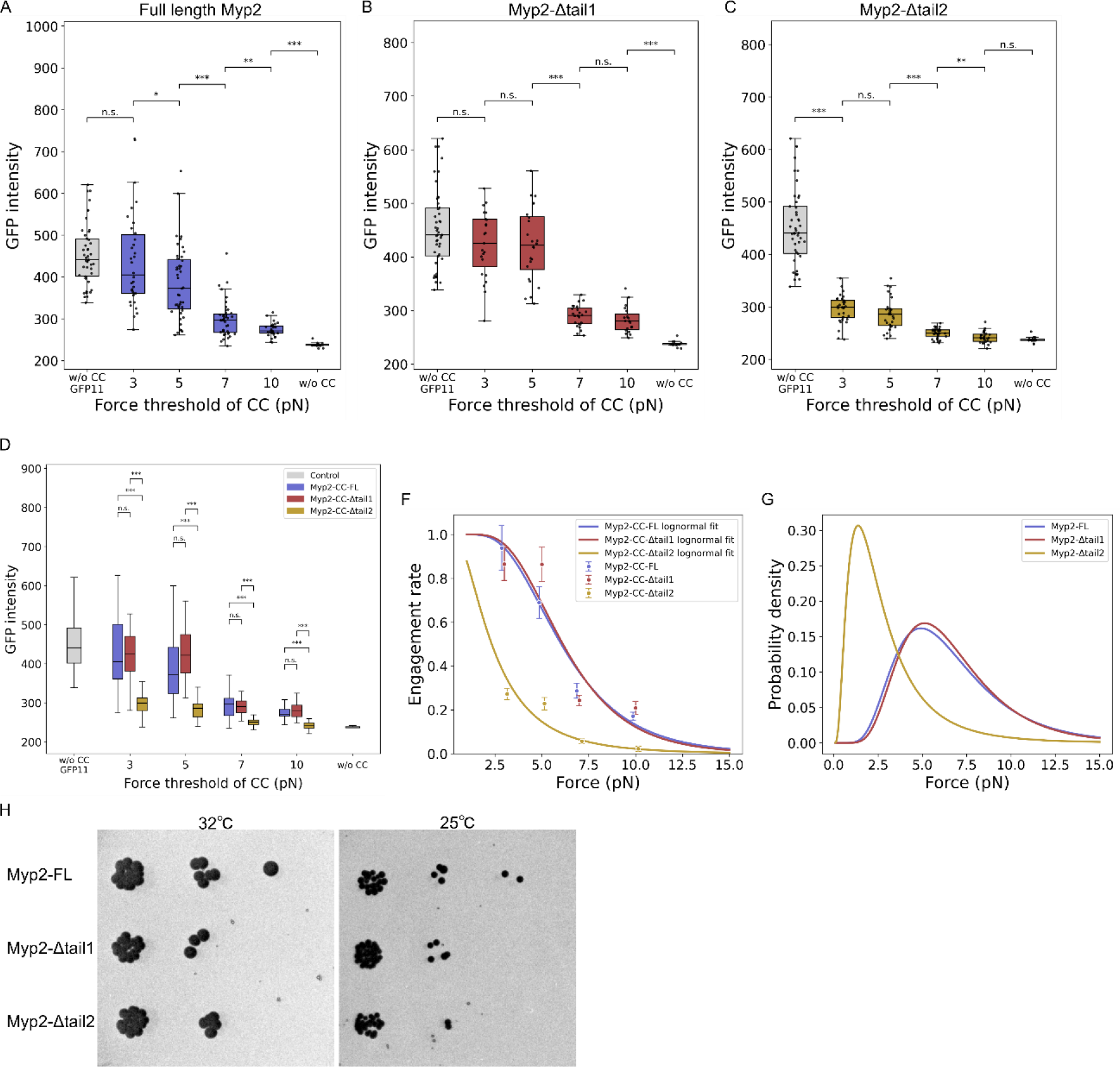
Quantitative analysis of CC sensor-based force measurements reveals distinct contributions of Myp2 tail domains. (A-D) Boxplots of GFP intensity in contractile rings in strains expressing (A) full-length Myp2 (Myp2-FL), (B) proximal tail deletion (Myp2-Δtail1), and (C) distal tail deletion (Myp2-Δtail2). (D) Comparison of GFP intensities between genotypes. (F) Engagement rates of strains expressing Myp2-FL (blue), Myp2-Δtail1 (red), and Myp2-Δtail2 (yellow) measured using CC sensors with force thresholds of 3, 5, 7, and 10 pN. Data were displayed as mean ± standard error. Solid lines represent regression curves of Eq. (2). (G) Probability density distributions of forces on Myp2-FL (blue), Myp2-Δtail1 (red), and Myp2-Δtail2 (yellow). (H) Strains expressing full-length Myp2, Myp2-Δtail1, and Myp2-Δtail2 grown on a YE5S plate at 32°C or 25°C for 4 days. Statistical significances were determined by One-way ANOVA with Tukey’s multiple comparison tests. ***, p < 0.001; **, p < 0.01; *, p < 0.05 and n.s., no significance. n > 14 rings per genotype.

In Myp2-Δtail2, no significant difference was detected between strains with CC-10pN and without CC, consistent with the colocalization analysis (Fig. 2L). In multiple comparisons between genotypes (Fig. 3D), no significant differences were detected between full-length Myp2 and Myp2-Δtail1 at any force threshold. In contrast, Myp2-Δtail2 exhibited statistically significant reductions in GFP intensity compared with both full-length Myp2 and Myp2-Δtail1 at all force thresholds.

By normalizing GFP intensity in contractile rings using intensity of strains with only GFP11 linker and without CC, we obtained engagement rates of each CC sensors defined as Eq. (1), *engagement rate* = (*I*_θ__cc_ − *I*_wo_)/(*I*_GFP11_ − *I*_wo_) (Fig. 3F). Due to the time-irreversible property of CC sensors, the engagement rate represents the cumulative force exposure of CC sensors with the threshold *θ*_cc_, corresponding to the probability density function of *P*(*F* > *θ*_cc_) in an appropriate probability distribution (Fig. S1). Given that the magnitude of the force is nonnegative due to thermal fluctuations, we used the log-normal distribution, *log F* ~ *N* (*μ, σ*^2^), where *μ* and *σ* are characteristic parameters.

Then, the engagement rate is rewritten by Eq. (2), 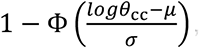, where Φ is the standard normal cumulative distribution function (CFD) (see Supplement). To determine the characteristic parameters of each genotype, we fit the linearized function of Eq. (2) to the experimentally obtained engagement rates using the least-square method (Fig. 3F). The resulting CDFs described the force probability distributions of full-length Myp2, Myp2-Δtail1, and Myp2-Δtail2 (Fig. 3G, Table S3). Compared with full-length Myp2 and Myp2-Δtail1, the tail2 deletion altered the force probability distribution. In log-normal distribution, the expected value (i.e. mean value) is obtained by 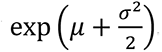. Full-length Myp2 and Myp2-Δtail1 r ues of 6.6 pN and 6.7 pN, respectively. In Myp2-Δtail2, the expected value was 3.0 pN, representing an approximately 55% reduction of the force level compared with full-length Myp2. This molecular-scale quantification of force is consistent with the reduced microscopic ring constriction rate observed upon deletion of the C-terminal tail ^14^. We also tested whether the tail deletion alters the growth rate. Myp2-Δtail1 and Myp2-Δtail2 exhibited slower growth than full- length Myp2 at 32°C and 25°C (Fig. 3H, S3). The deletion of tail2 more clearly slowed growth at 25°C, consistent with the previous results of growth assay at the temperature ^14^. Taken together, the C-terminal tail of Myp2 is required for efficient force transmission across the scale.

### The C-terminal tail of Myp2 alone is sufficient for clustering but not for ring localization

Live-cell imaging has reported that Myp2 appears as motile clusters on the ring, indicating a spatially heterogeneous organization rather than a continuous distribution ^19^. Considering the robust function of Myp2, this motile cluster may locally rescue stability and efficient constriction of the ring. In contrast, at the molecular scale, Myp2 exhibits slower molecular turnover than Myo2 ^20^, meaning that Myp2 is immobile. This immobility is caused by the C-terminal tail region of Myp2 ^14^. These studies attribute the formation of motile clusters to the C-terminal tail of Myp2. Thus, we tested whether the C-terminal tail itself is involved in the formation of clusters in cells. Myp2 tail2 tagged with mNeonGreen2 was inserted after a self-cleavage 2A peptide at the C-terminal end of *hxk2* gene, which is expressed in very high copy number ^21^. The over-expression of tail2 in the Myp2-Δtail2 strain formed huge clusters in the cytoplasm (Fig. 4A). In contrast, the over-expression of tail2 in the strain expressing full-length Myp2 exhibited the localization of the clusters to the ring (Fig. 4B). These data indicate that the tail2 region itself is not sufficient for the localization of clusters to contractile rings.

**Figure 4.**
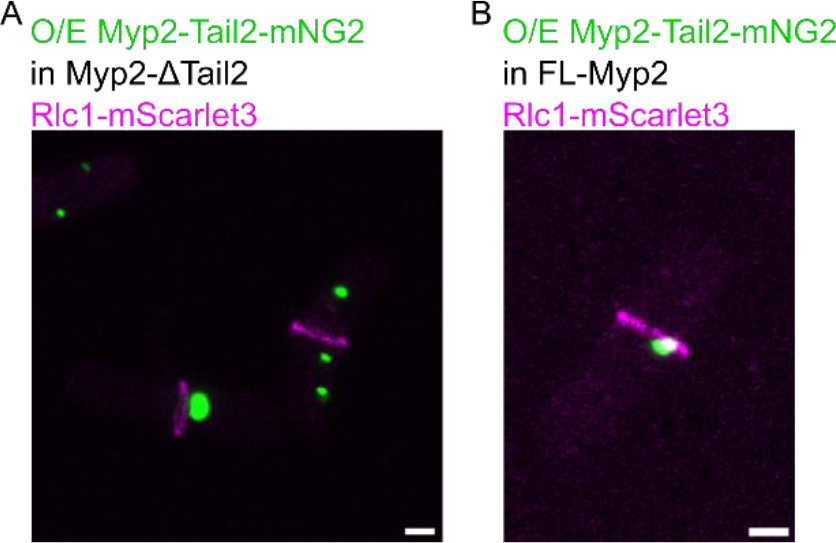
Myp2 tail2 forms clusters by self-oligomerization but does not localize to contractile rings by itself. (A) Over-expression of Myp2-tail2 tagged with mNeonGreen2 in a strain expressing Myp2-Δtail2 and Rlc1-mScarlet3. (B) Over-expression of Myp2-tail2 tagged with mNeonGreen2 in a strain expressing full-length Myp2 and Rlc1-mScarlet3. Scale bar; 2 μm.

### Discussion

Fission yeast *S. Pombe* has two type-II myosins, Myo2 and Myp2. Myo2 contributes to the contractile ring assembly at the early stage of cytokinesis, whereas unconventional myosin Myp2 is crucial to the ring constriction ^13,19,22,23^. Under stress conditions such as high osmotic pressure or when Myo2 is suppressed, Myp2 plays a more important role in the ring constriction or even in the ring assembly to rescue Myo2 ^13,14^. Unlike the canonical tail structure of Myo2 and other myosin-II, Myp2 possesses two distinct coiled-coil regions connected by a nonhelical hinge, folding back and forming an intramolecular antiparallel coiled-coil *in vitro* ^11,15^. This unique tail architecture may contribute to the robust function of Myp2 *in vivo*. Although these studies have revealed the molecular function and regulation of Myp2, it is still challenging to directly and quantitatively determine the mechanistic contribution of Myp2 and its tails to contractile rings at the molecular and intramolecular scale. To this end, we measured forces on Myp2 and the tail deletion mutants using recently developed coiled-coil force sensors ^16,17^. Combining force measurements and quantitative analysis, we showed that the C-terminal tail region (referred to as tail2 in this study), but not the N-terminal tail, predominantly contributes to load-bearing of Myp2.

Forces of myosin-II motors in theoretical models have typically been estimated based on single-molecule measurements of muscle myosin, which report forces of a few piconewtons per motor head ^24–27^. These values obtained by muscle myosin have been widely adopted in computational models of the contractile ring ^28–31,3,32^. Although FRET-based tension sensors inserted in non-muscle myosin-II revealed spatial and temporal heterogeneity of FRET-index in living cells ^33^, force quantification of non-muscle myosin-II in *vivo* has been technically challenging due to its complicated domain and molecular organizations. In this study, CC force sensors and quantitative analysis revealed that Myp2 bears 6.6 pN of the mean value in living cells, which is consistent with forces measured for muscle myosin and thus supports the above previous computational models. We also found that Myp2-Δtail2 bears 3.0 pN, showing approximately 55% reduction compared to full-length Myp2 and Myp2- Δ tail1. This data indicates the mechanical contribution of tail2 *in vivo*. In addition to the mean value of forces, we obtained the force probability distributions of full-length Myp2, Myp2-Δtail1, and Myp2-Δtail2 (Fig. 3G, Table S3), providing more quantitative constraints for theoretical models.

The folded conformation of myosin-II tails is evolutionarily conserved across eukaryotes, where it typically functions as an autoinhibition. Myosin-II molecules have been shown to adopt a folded, autoinhibited conformation that can unfold into an active state upon regulatory signals such as phosphorylation ^34^. Activation of the non-muscle myosin II promotes its assembly into bipolar filaments, which are essential for the formation, maturation, and maintenance of contractile actomyosin structures such as stress fibers ^35–37,32,38^. In fission yeast, Myp2 is a monomeric myosin, forming the antiparallel folded back tail structure with C-terminal and N-terminal coiled-coils *in vitro* ^15^. In living fission yeast, this tail structure, particularly the C-terminal tail region, is important for the robust ring constriction upon stress conditions during cytokinesis *in vivo* ^14^, forming spatially heterogenous motile clusters on the ring ^19^. The over-expression of the tail2 region exhibited huge clusters and localized to the ring in the presence of full-length Myp2 (Fig. 4A, 4B), indicating that these motile clusters of Myp2 are formed by the self-oligomerization of tail2. These observations suggest that the C-terminal tail contributes to higher-order organization of Myp2, which may in turn influence the mechanical forces borne by the molecule. Considering the reduction of force on Myp2 caused by the tail2 deletion (Fig. 3G), together with our observation and the previous findings of Myp2 ^15,19,14^, the folded back structure of the tail region may be mechanically engaged *in vivo*.

## Material and Methods

### Yeast strains and cell culture

*Schizosaccharomyces pombe* strains were maintained in YE5S composed of yeast extract supplemented with adenine, histidine, leucine, lysine, and uracil at 0.225 g/L. Strains used in this study were constructed using the CRISPR-Cas9 genome editing technique with appropriate primers (See Supplement) ^39,40^.

### Coiled-coil force sensors

Coiled-coil (CC) force sensors composed of anti-parallel coiled-coil peptides connected with a GFP11 linker were used for force measurements on Myp2 ^16,18^. CC force sensors with force thresholds of 3, 5, 7, and 10 pN (referred to as CC-3pN, CC-5pN, CC-7pN, and CC-10pN, respectively) were genetically inserted *myp2* gene. Non-fluorescent GFP1-10 was expressed in cytoplasm by replacing *pil1* gene.

### Growth assay

Fission yeast strains were cultured in YE5S until OD_600_∼0.5 and resuspended by water to adjust OD_600_ = 0.1. Then, 5 µL drops of each strain with 10^2^, 10^3^, and 10^4^ dilutions were placed on a YE5S agarose plate. The plates were maintained for 4 days in 32°C or 25°C incubator. Images of the plates growing colonies were acquired by a gel imager. An inverted LUT in ImageJ/Fiji was used for visualization of these images.

### Microscopy

A glass-bottom dish was treated by 50 µg/mL lectin (Sigma, L2380) and dried in a 32°C incubator. Fission yeast strains grown until an optical density OD_600_=0.3-0.6 were placed onto the lectin-treated dish. Fluorescence images were sequentially acquired every 0.5 µm for 10 µm height (21 optical z-sections) with a 60x/NA1.42 objective mounted on a laser scanning confocal microscope FV3000.

### Quantitative analysis

Maximum intensity projection was performed using ImageJ/Fiji. For colocalization analysis to distinguish between open and closed states of CC sensors with split -GFP readout, Pearson’s correlation coefficient (R-value) was obtained by pixel-wise fluorescence intensity of ring marker Rlc1 tagged with mScarlet3 and GFP of CC sensors within a cell.

For a quantitative analysis of force probability distribution, contractile rings were identified by Rlc1-mScarlet3. An engagement rate representing the fraction of molecules experienced forces exceeding a coiled-coil force threshold is defined as

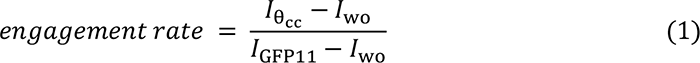

where *I*_θ__cc_, *I*_wo_, and *I*_GFP11_ represent green fluorescence intensity of rings in strains expressing Myp2 with a CC sensor of an opening force threshold *θ*_cc_, GFP11, and no insertion, respectively. Because CC force sensors are fluorescent by forces exceeding the opening thresholds, the engagement rate reflects a probability density function (PDF) represented by *P*(*F* > *θ*_cc_) with an appropriate probability distribution (Fig. S1). Due to the presence of thermal fluctuations, forces on Myp2 are nonnegative. Therefore, we assumed a log-normal distribution of force *F* on Myp2, i.e.

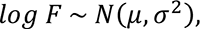

an analytical engagement rate is described by

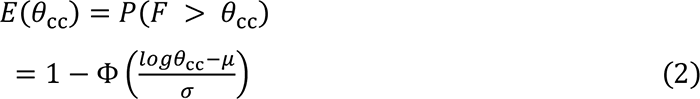

where Φ denotes the standard normal cumulative distribution function (CDF) (see Supplement). To determine *μ* and *σ*, the engagement rate was fitted by a linearized regression curve with the least-square method (See supplement):

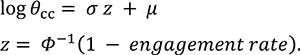

## Statistical analysis

Results are displayed as mean ± standard deviation unless otherwise notified. Statistical significance was determined by one-way ANOVA with Tukey’s multiple comparison tests for multiple groups. For Pearson’s correlation coefficient (R-value), statistical significance was determined by Permutation tests. ***, p < 0.001; **, p < 0.01; *, p < 0.05 and n.s., no significance.

## Acknowledgements

We thank Dr. Bertocchi, Dr. Daiki Matsunaga, and Naoto Yonekura for helpful comments, Dr. Yuan Ren for his pioneering work for coiled-coil sensors, Andras Bencze for inspiring discussions, Tsukasa Inoue for technical support, and TOKUSHUKAI Scholarship for supporting J.L.

## Funding

This study was partly supported by JSPS (Japan Society for the Promotion of Science) KAKENHI 25K21545 and 26H00969.

## Author contributions

Conceptualization: J.L. and T.S. Methodology: T.S. and J.B. Investigation: J.L. and T.S. Visualization: J.L. Funding acquisition: T.S. Project administration and Supervision: T.S. Writing—original draft: J.L. Writing—review and editing: T.S., S.D., and J.B.

## Competing interests

Julien Berro has a patent US20250368701A1 regarding coiled-coil force sensors used in this study. Other authors declare no competing financial interests or personal relationships that could influence this study.

## Data and materials availability

All data are present in the main text and/or the supplementary materials. The strains and plasmids are available from Julien Berro or Takumi Saito upon reasonable requests.

