## Supplementary material for "Mechanical contributions of the Myp2 tail revealed by coiled-coil force sensors": Fig. S1

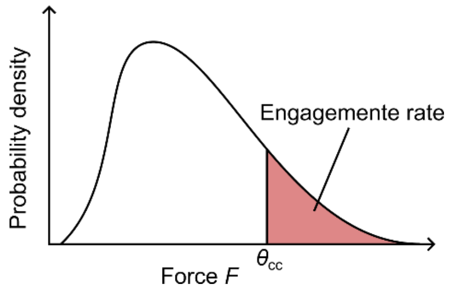


Figure S1 Schematic illustration of the engagement rate corresponding to a probability density function ($P(F > \theta_{\mathrm{cc}})$) in an appropriate probability distribution.
