## Supplementary material for "Mechanical contributions of the Myp2 tail revealed by coiled-coil force sensors": Fig. S2

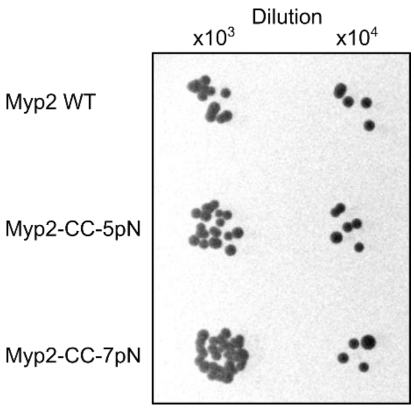


Figure S2 Strains expressing Myp2 with and without coiled-coil sensors were grown on a YE5S plate at 32℃ for 4 days.
