## Supplementary material for "Mechanical contributions of the Myp2 tail revealed by coiled-coil force sensors": Fig. S3

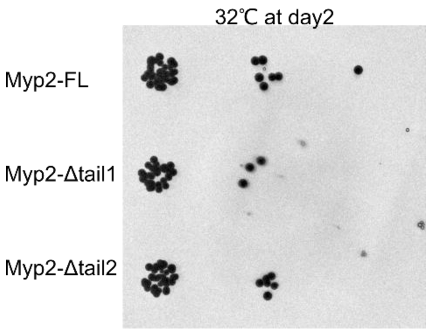


Figure S3 Strains expressing full-length Myp2, Myp2-Δtail1, and Myp2-Δtail2 were grown on a YE5S plate at 32℃ for 2 days.
