## Supplementary material for "Mechanical contributions of the Myp2 tail revealed by coiled-coil force sensors": Table S1

Table S1 The list of fission yeast strains used and constructed in this study.

| SpTS | Genotype |
|---|---|
| 14 | rlc1-mScarlet3, GFP1-10 replaces pil1 fex1Δ fex2Δ ade6-M216 his3-D1 leu1-32 ura4-D18 |
| 21 | myp2-Q833-DyneinStalk-GFP11, rlc1-mScarlet3, GFP1-10 replaces pil1 fex1Δ fex2Δ ade6-M216 his3-D1 leu1-32 ura4-D18 |
| 23 | myp2-Q833-Oakley-p1-LV-L3S-GFP11, rlc1-mScarlet3, GFP1-10 replaces pil1 fex1Δ fex2Δ ade6-M216 his3-D1 leu1-32 ura4-D18 |
| 24 | myp2-Q833-Oakley-p1-GFP11, rlc1-mScarlet3, GFP1-10 replaces pil1 fex1Δ fex2Δ ade6-M216 his3-D1 leu1-32 ura4-D18 |
| 25 | myp2-Q833-Oakley-p1-OnePlus-GFP11, rlc1-mScarlet3, GFP1-10 replaces pil1 fex1Δ fex2Δ ade6-M216 his3-D1 leu1-32 ura4-D18 |
| 33 | myp2-Q833-Oakley-p1-LV-L3S-GFP11-delta1611-end, rlc1-mScarlet3, GFP1-10 replaces pil1 fex1Δ fex2Δ ade6-M216 his3-D1 leu1-32 ura4-D18 |
| 34 | myp2-Q833-Oakley-p1-GFP11-delta1611-end, rlc1-mScarlet3, GFP1-10 replaces pil1 fex1Δ fex2Δ ade6-M216 his3-D1 leu1-32 ura4-D18 |
| 34 | myp2-Q833-Oakley-p1-oneplus-GFP11-delta1611-end, rlc1-mScarlet3, GFP1-10 replaces pil1 fex1Δ fex2Δ ade6-M216 his3-D1 leu1-32 ura4-D18 |
| 36 | myp2-delta1611-end, rlc1-mScarlet3, GFP1-10 replaces pil1 fex1Δ fex2Δ ade6-M216 his3-D1 leu1-32 ura4-D18 |
| 41 | myp2-Q833-GFP11, rlc1-mScarlet3, GFP1-10 replaces pil1 fex1Δ fex2Δ ade6-M216 his3-D1 leu1-32 ura4-D18 |
| 45 | myp2-Q833-DyneinStalk-GFP11-delta1611-end, rlc1-mScarlet3, GFP1-10 replaces pil1 fex1Δ fex2Δ ade6-M216 his3-D1 leu1-32 ura4-D18 |
| 51 | myp2-delta834-1248, rlc1-mScarlet3, GFP1-10 replaces pil1 fex1Δ fex2Δ ade6-M216 his3-D1 leu1-32 ura4-D18 |
| 52 | myp2-Q833-DyneinStalk-GFP11-delta834-1248, rlc1-mScarlet3, GFP1-10 replaces pil1 fex1Δ fex2Δ ade6-M216 his3-D1 leu1-32 ura4-D18 |
| 53 | myp2-Q833-Oakley-p1-LV-L3S-GFP11-delta834-1248, rlc1-mScarlet3, GFP1-10 replaces pil1 fex1Δ fex2Δ ade6-M216 his3-D1 leu1-32 ura4-D18 |
| 54 | myp2-Q833-Oakley-p1-GFP11-delta834-1248, rlc1-mScarlet3, GFP1-10 replaces pil1 fex1Δ fex2Δ ade6-M216 his3-D1 leu1-32 ura4-D18 |
| 55 | myp2-Q833-Oakley-p1-oneplus-GFP11-delta834-1248, rlc1-mScarlet3, GFP1-10 replaces pil1 fex1Δ fex2Δ ade6-M216 his3-D1 leu1-32 ura4-D18 |
| 63 | hxk2-ERBV-myp2-tail2-mNeonGreen2, myp2-delta1611-end, rlc1-mScarlet3, GFP1-10 replaces pil1 fex1Δ fex2Δ ade6-M216 his3-D1 leu1-32 ura4-D18 |
