## Supplementary material for "Mechanical contributions of the Myp2 tail revealed by coiled-coil force sensors": Table S2

Table S2 The list of primers used in this study.

| TS' | Sequence |
|---|---|
| 40 | AGGATGTGGCTTCAATAACTttcTTCGGTACAGGTTATG |
| 41 | AGTTATTGAAGCCACATCCTGTTTTAGAGCTAGAAATAGCAAG |
| 42 | GGCAACAATAACGATCATCAA |
| 43 | GCCAGAGCTTGTTCCTCTT |
| 44 | CCCGGCTATTGGCTCTTT |
| 51 | GCCAGTGTCTTGGGTTCTT |
| 52 | TAGAGAGGGAATACAGATTGTTCTTCGGTACAGGTTATG |
| 53 | CAATCTGTATTCCCTCTCTAGTTTTAGAGCTAGAAATAGCAAG |
| 54 | CGGTCAGTTACACAACACACTT |
| 55 | CGGGGAAGGTCTGTCAGAAT |
| 56 | CAGCTTCAGAAGGAAAGGTGTA |
| 96 | TCGCAAGCTGTTCGCGGTACTTCTTCGGTACAGGTTATG |
| 97 | GTACCGCGAACAGCTTGCGAGTTTTAGAGCTAGAAATAGCAAG |
| 98 | GGGTTGCAAGGACCTTCTGT |
| 99 | CGCCGCCTATCTCTTCATA |
| 100 | GGCAAAATGGAAGCACTGAT |
| 124 | GCGGAGGACGAGACCATA |
| 164 | CCATCAATGCGTTTTAGAGGAT |
| 165 | GACAAGACGGAGACTTTACCGGTTTAATCATAGGCAAGAT |
| 174 | TTGCAACCACTCGGTCATGGTGGCGACCGGTAG |
| 175 | ACCGAGTGGTTGCAAGAAGAA |
| 176 | ATTAATAAGACCAGGAATACGCCTTAGAACGCTAGGCGAAC |
| 177 | CACGTTCTTCTTCCAACTGTACT |
| 207 | TCTCTTCTTATTTACCCACATCTATTTCTATTTCTGCAAGATGACCGAGTGGTTGCAAGA |
