## Supplementary material for "Mechanical contributions of the Myp2 tail revealed by coiled-coil force sensors": Table S3

Table S3 Characteristic parameters for log-normal distributions of full-length Myp2, Myp2-Δtail1, and Myp2-Δtail2

| | $\mu$ | $\sigma$ |
|---|---|---|
| Full-length Myp2 | 1.79 | 0.45 |
| Myp2-Δtail1 | 1.81 | 0.42 |
| Myp2-Δtail2 | 0.84 | 0.72 |
