## Supplement for "Mechanical contributions of the Myp2 tail revealed by coiled-coil force sensors"

**Linearized regression curve of the engagement rate to determine a force probability distribution.**

We fit the measured engagement rate to the analytical engagement rate

$$engagement\ rate = E(\theta_{\mathrm{cc}}).$$

Thus,

$$engagement\ rate = 1 - \Phi\left(\frac{(log\ \theta_{\mathrm{cc}} - \mu)}{\sigma}\right)$$

where $\Phi$ is the standard normal cumulative distribution function (CDF) defined as

$$\Phi(\mathrm{z}) = \frac{1}{\sqrt{2\pi}} \int_{-\infty}^{z} e^{-\frac{t^2}{2}} dt.$$

By taking inverse of the CDF,

$$log\ \theta_{\mathrm{cc}} - \mu = \sigma\, \Phi^{-1}(1 - engagement\ rate)$$

the linearized regression curve

$$\log \theta_{\mathrm{cc}} = \sigma z + \mu$$
$$z = \Phi^{-1}(1 - engagement\ rate)$$

is obtained.
